# Dix-seq: An integrated pipeline for fast amplicon data analysis

**DOI:** 10.1101/2020.05.11.089748

**Authors:** Yongjun wei, Tianqi Ren, Lei Zhang

**Author notes:** Corresponding authors, Yongjun Wei, Lei Zhang.

## Abstract

The amplicon derived from 16S rRNA genes, 18S rRNA genes, internal transcribed spacer sequences or other functional genes can be used to infer and evaluate microbial diversity or functional gene diversity. With the development of sequencing technologies, large amounts of amplicon data were generated. Several different software or pipelines had been developed for amplicon data analyses. However, most current software/pipelines require multistep and advanced programming skills. Moreover, they are often complex and time-consuming. Here, we introduced an integrated pipeline named Dix-seq for high-throughput amplicon sequence data processing (https://github.com/jameslz/dix-seq). Dix-seq integrates several different amplicon analysis algorithms and software for diversity analyses of multiple samples. Dix-seq analyzes amplicon sequences efficiently, and exports abundant visual results automatically with only one command in Linux environment. In summary, Dix-seq enables the common/advanced users to generate amplicon analysis results easily and offers a versatile and convenient tool for researchers.

## 1. Introduction

The microbial diversity and functional information can be inferred with the amplicon sequencing generated from marker genes, such as 16S rRNA gene, 18S rRNA gene, internal transcribed spacer sequences, and other functional genes. The low cost of next-generation sequencing technology and popularization of amplicon study, as well as the easy application of amplicon technology, lead to the generation of large amount of amplicon data. To show and compare microbial communities, several different amplicon analysis tools and pipelines have been developed. These tools or pipelines often require the users to master in-depth bioinformatics knowledge and understanding of parameters in each step during amplicon analyses. Moreover, the obtained output needs to be visualized with additional software/tools. Besides the requirement of computer professional knowledge, the whole amplicon analysis process is complex and time-consuming. Here, we develop an integrated pipeline named Dix-seq for convenient amplicon data analyses (https://github.com/jameslz/dix-seq). Dix-seq generates abundant visual results automatically with only one command line in Linux environment, from the raw data demultiplex, chimeric sequence removal, sequence classification, alpha and beta diversity, functional prediction, variance analyses, to the final visual results (Figure 1). The pipeline is easy and efficient for further improvement, for diverse third-party procedures can be integrated and pipeline optimization are allowed (Table 1).

**Figure 1.**
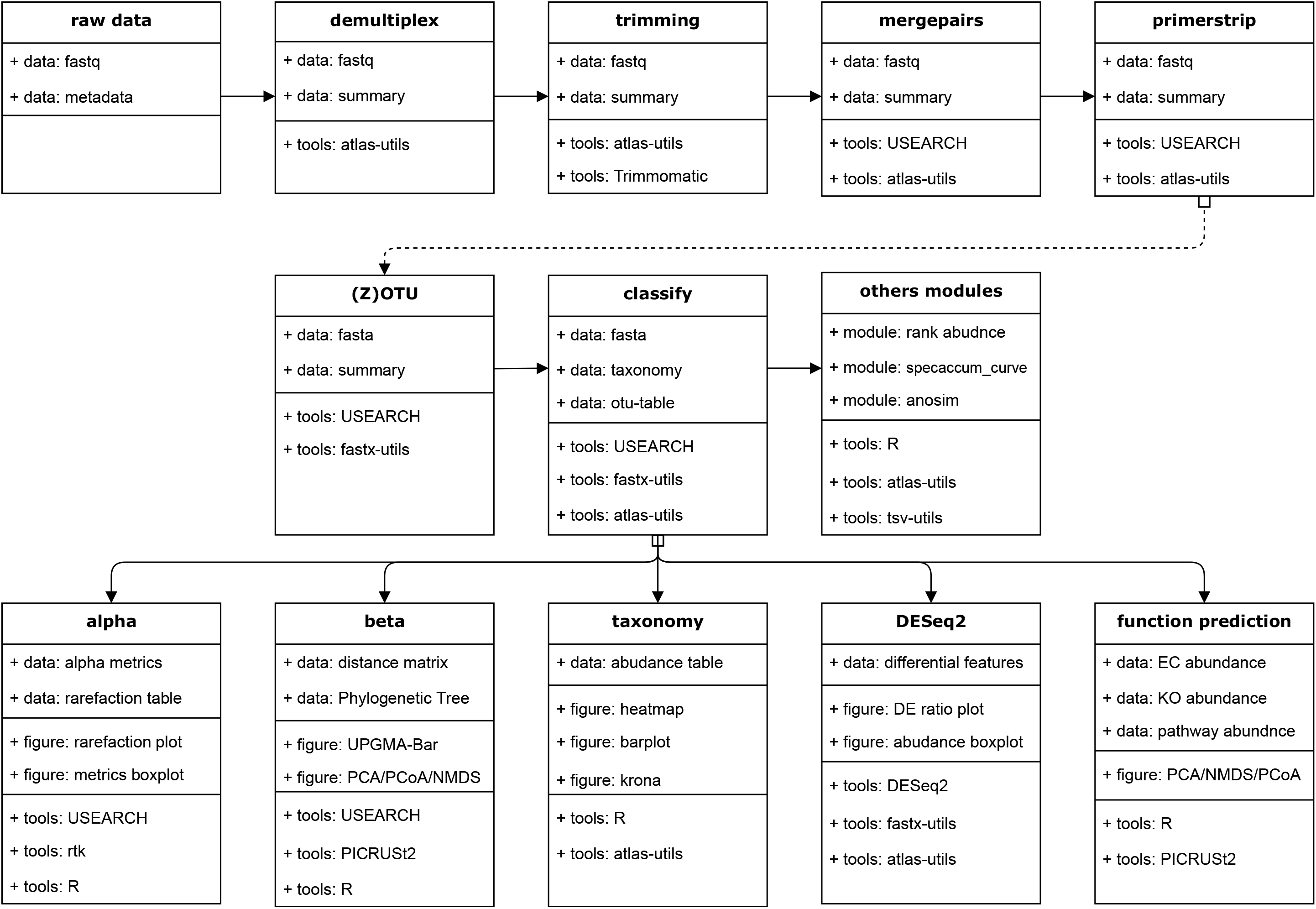
Dix-seq: An integrated pipeline for fast amplicon data analysis. The steps and softwares used in the Dix-seq pipeline were shown.

**Table 1.**
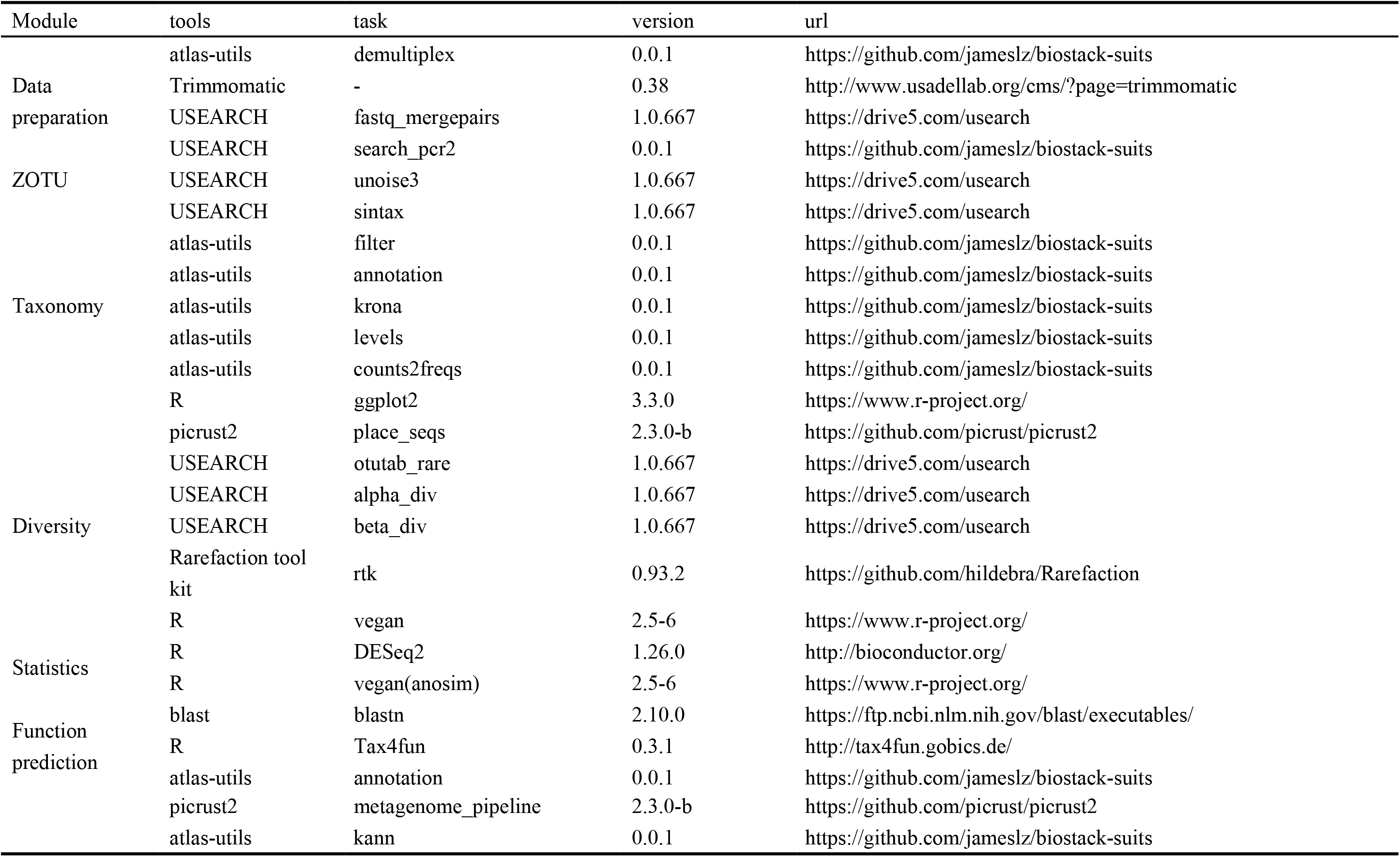
Details of softwares and tools used in Dix-seq pipeline.

## 2. Raw sequence data analyses

Dix-seq can extract raw fastq files directly from the sequencing machines in short time. Next, raw fastq files are demultiplexed with the corresponding barcode sequences by our developed atlas-utils tool (https://github.com/jameslz/biostack-suits version: 0.0.1). After discarding barcode sequences in each read, low-quality sequences were filtered by Trimmomatic (version 0.38) [1]. During filter process, the start and end of a read below a threshold quality of score < 3 were removed, and the reads were truncated at any site receiving an average quality score < 15 over a 4 bp sliding window. Moreover, truncated reads which were shorter than 36bp were discarded. Paired-end reads were merged using Usearch fastq_mergepairs command with the default parameters (usearch11.0.667, http://drive5.com) [2]. Reads were discarded if they could not be merged. The primer sequences would affect further operational taxonomic unit (OTU) table construction. Therefore, merged reads without full primer sequences or with more than two nucleotide mismatches in the primer region were discarded, and the clean merged reads (clean tags) without primer sequences were obtained.

## 3. Bioinformatic analyses

### 3.1 ZOTU Construction

In the pipeline, Usearch fastq_filter was used to filter all the low-quality sequences in the clean tags with default parameters (fastq_maxee 1), and the high-quality sequences of the clean tags were obtained. The obtained high-quality reads were sorted in non-redundant abundance order using USEARCH fastx_uniques. The USEARCH unoise3 algorithm filters the non-redundant sequences with minsize = 8, and generate the zero-radius operational taxonomic unit (ZOTU) table with 97% identity. The representative sequences of each ZOTU was identified. During the process, the chimeric sequences were removed in the denoising procedure. All the high-quality sequences were binned to the ZOTU representative sequences, and the relative abundance of each ZOTU were obtained.

The phylogenetic assignment of the representative sequences of each ZOTU was determined by **Usearch sintax** algorithm [4] with the RDP training set (version **v16**) 16S rRNA database (Other functional gene database can be used, such as nifH sequence database) as reference database (the confidence threshold is 0.8). The ZOTUs annotated as mitochondrial or chloroplast rRNA sequences were discarded. As a result, ZOTU counts table and ZOTU relative abundance table were generated. The **stacked bar charts** and **KRONA** software (http://sourceforge.net/projects/krona/) were used for visualization of the taxonomic composition and phylogenetic annotation. The heatmap plots at different taxonomic levels (ZOTU, genus, family, order, class, phylum) were generated using**pheatmap** in R environment (version 3.6.0) [5]. In the meanwhile, the clustering analyses between different samples or ZOTUs were applied.

The ZOTU table was normalized using USEAERCH otutab_rare with specific sample size, and the normalized sample size was the sample harboring the minimum sequences among all the samples. The diversity and other analyses were based on the normalized ZOTU table.

### 3.2 Diversity and functional analyses

Based on ZOTU abundance table, the alpha diversity indices of richness, chao1, simpson and shannon diversity were calculated with Usearch alpha_div. The ZOTU representative sequences were aligned with the reference sequences using HMMER and EPA-NG method in PICRUSt2 (version 2.3.0-b), and the multiple alignment results was used to generate the phylogenetic tree.

The beta diversity distance was calculated with **Usearch beta_div**, and the frequently used distance metrics were weighted_unifrac, unweighted_unifrac and bray_curtis. The principal coordinates analysis (PCoA) and Non-metric Multidimensional Scaling (NMDS) analyses based on weighted_unifrac, unweighted_unifrac and bray_curtis distances were used to evaluate the variation/difference between samples based on the homogenized OTU table. The gmodels in R software and nmds.py were used to generate the final figures PCoA and NMDS figures. For unweighted and weighted UniFrac distance metrics, the required phylogenetic tree was generated using **PICRUSt2 in previous step**.

The **Vegan** 2.4.2 package [6] in R was used to generate Anosim based on the distances calculated with homogenized OTU tables. The clean tags were used for differential abundance analysis at different taxonomic levels (phylum, order, class, family, genus, ZOTU) using **DEseq2 with the parameters of test=Wald and fitType=parametric**.

The R package **Tax4Fun** (version 0.3.1) [6] was used to predict the functional profiles with the information of ZOTU representative sequences and their relative abundance. During the process, **SILVA_123_SSURef_Nr99** database in Tax4Fun was used to identify the best hit for representative ZOTU sequences, and the predicted KO (KEGG Orthology) abundance was used to infer pathway, enzyme, module and catalog information in KEGG database. The predicted functional information of samples was compared and shown.

#### Demo amplicon analyses

Dix-seq was used to analyze one amplicon data with six samples (Accession number of PRJNA436765). The amplicon sequences were V3-V4 fragments of 16S rRNA genes, and the six samples were divided into three groups. Dix-seq generates visual results in several minutes. Part of the visual outputs were shown in Figure 2. The phylum-level microbial abundance and the phylogenetic classification of these six samples can be generated (Figure 2a), and other phylogenetic level abundance results can also be generated. The alpha diversity indices, including richness, were calculated (Figure 2b). Based on the principal components, the relationship between these six samples were analyzed (Figure 2c). Moreover, other microbial community differences, such as the unweighted pair group method with arithmetic mean (UPGMA), were analyzed (Figure 2d). Other beta diversity distances were visualized with Dix-seq. Principal component analysis diagram shows microbial composition differences of the six samples based on prediction of Tax4Fun at pathway-level was expressed with the principal component analysis (Figure 2e), and other functional levels such as KO, modules and catalog, were predicted and expressed. In addition, the pair-wise comparison between two groups at OTU-level was calculated with Deseq (Figure 2f), other phylogenetic level and variance between each two group were analyzed.

**Figure 2.**
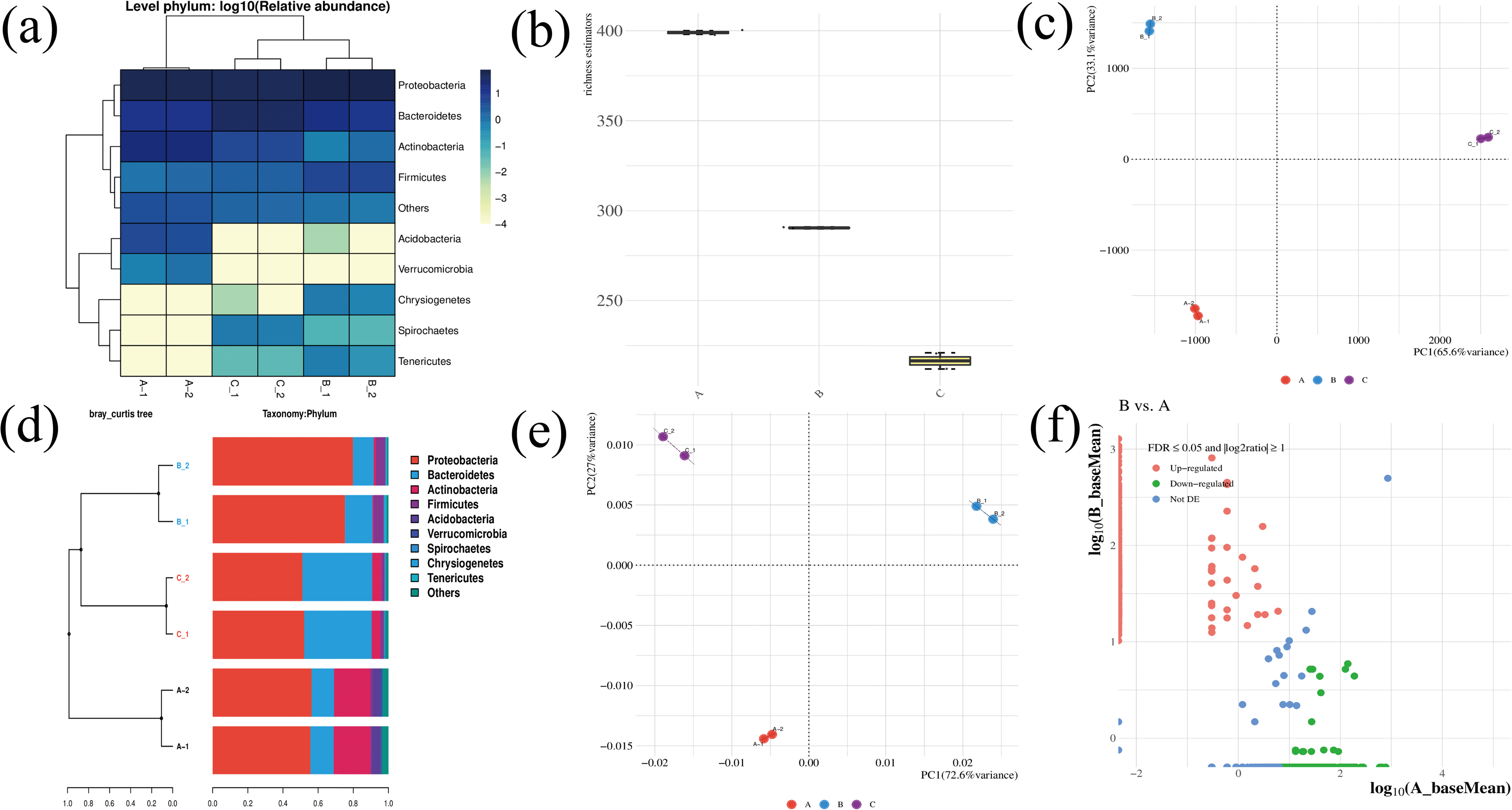
Some outputs generated with the Dix-seq pipeline were shown. (a) Microbial abundance of six samples at phylum-level. (b) The richness estimation of the six samples. (c) Principal component analysis diagram shows microbial composition differences of the six samples. (d) UPGMA diagram shows microbial composition differences of the six samples. The microbial compositions at phylum-level of the six samples are shown. (e) Principal component analysis diagram shows microbial composition differences of the six samples based on prediction of Tax4Fun at pathway-level. (f) Pair-wise comparison in Deseq at ZOTU-level between Group_A and Group_B.

## 4. Conclusion

We developed an integrated pipeline for efficient amplicon data analyses. There are several advantages of this pipeline. Firstly, the developed Dix-seq pipeline is based on command line model, and all the visual output can be easily realized with one command line. Secondly, Dix-seq pipeline is composed with different functional workflow modules, and these modules can work together or separately. Moreover, the functional modules are independent PERL scripts and can be added or deleted easily. Thirdly, the pipeline integrates three commond–line tools: atlas-utils, tsv-utils, fastx-utils (packed in biostack-suits), which can generate results efficiently. Finally, Dix-seq pipeline integrates several third-party procedures, including functional prediction, variance analysis. In summary, our integrated Dix-seq pipeline offers a versatile and efficient amplicon analysis tool for different-level researchers.

